# Optimizing genomic prediction of host resistance to koi herpesvirus disease in carp

**DOI:** 10.1101/609784

**Authors:** Christos Palaiokostas, Tomas Vesely, Martin Kocour, Martin Prchal, Dagmar Pokorova, Veronika Piackova, Lubomir Pojezdal, Ross D. Houston

## Abstract

Genomic selection (GS) is increasingly applied in breeding programmes of major aquaculture species, enabling improved prediction accuracy and genetic gain compared to pedigree-based approaches. Koi Herpesvirus disease (KHVD) is notifiable by the World Organisation for Animal Health and the European Union, causing major economic losses to carp production. Genomic selection has potential to breed carp with improved resistance to KHVD, thereby contributing to disease control. In the current study, Restriction-site Associated DNA sequencing (RAD-seq) was applied on a population of 1,425 common carp juveniles which had been challenged with Koi herpes virus, followed by sampling of survivors and mortalities. Genomic selection (GS) was tested on a wide range of scenarios by varying both SNP densities and the genetic relationships between training and validation sets. The accuracy of correctly identifying KHVD resistant animals using genomic selection was between 8 and 18 % higher than pedigree best linear unbiased predictor (pBLUP) depending on the tested scenario. Furthermore, minor decreases in prediction accuracy were observed with decreased SNP density. However, the genetic relationship between the training and validation sets was a key factor in the efficacy of genomic prediction of KHVD resistance in carp, with substantially lower prediction accuracy when the relationships between the training and validation sets did not contain close relatives.

## Introduction

Genomic selection has become a cornerstone of genetic improvement in both plant and livestock breeding, enabling improved prediction accuracy, control of inbreeding, and (in some cases) reduction in generation interval compared to traditional pedigree-based approaches (Meuwissen *et al.* 2016; Hickey *et al.* 2017). The landmark paper of Meuwissen *et al.* (2001) highlighted the concept of breeding value prediction based on the joint merit of all markers distributed throughout the genome, and the advent of high-throughput DNA sequencing and development of SNP arrays in the subsequent decade made this concept a practical reality. While the application of genomics in aquaculture breeding has traditionally lagged behind the plant and terrestrial livestock sector, it is gaining momentum with reference genome assemblies and SNP arrays now available for most of the key aquaculture species (Yue and Wang 2017; Robledo *et al.* 2017). Both simulation and empirical studies suggest that considerable improvement in breeding value prediction accuracy is plausible, even with relatively modest SNP marker densities (Sonesson and Meuwissen 2009; Lillehammer *et al.* 2013; Ødegård *et al.* 2014; Tsai *et al.* 2015; Vallejo *et al.* 2018; Vallejo *et al.* 2017; Correa *et al.* 2017; Robledo *et al.* 2018).

Infectious diseases present a major and persistent threat to sustainable aquaculture production, and breeding for improved host resistance is an increasingly important component of mitigation (Houston 2017). Common carp *(Cyprinus carpio)* is one of the world’s most important freshwater aquaculture species, particularly in Asia and Europe. However, koi herpesvirus disease (KHVD), also known as Cyprinid herpesvirus-3 (CyHV-3) disease is a major threat to carp farming and is listed as a notifiable disease by the European Union (Taylor *et al.* 2010) and the World Organization for Animal Health (OIE 2018). Encouragingly, resistance to KHVD has been shown to be a highly heritable trait with estimates ranging between 0.50 – 0.79 (Ødegård *et al.* 2010; Palaiokostas *et al* 2018a). The potential of selective breeding for improved KHVD resistance in carp (utilizing information from challenge trials) has been illustrated by several studies which demonstrated large variation in survival both between-family (Dixon *et al.* 2009; Tadmor-Levi *et al.* 2017) and between strain (Shapira *et al.* 2005; Piačková *et al.* 2013). Further, a significant QTL associated with resistance to KHVD has been identified (Palaiokostas *et al.* 2018a). Nevertheless, the potential of genomic selection for improving KHVD resistance in carp has not yet been studied.

While SNP arrays are available for several aquaculture species, and are commonly used in some of the most advanced commercial breeding programmes (e.g. Atlantic salmon), they tend to be relatively expensive and can suffer from ascertainment bias (Robledo *et al.* 2017). Genotyping by sequencing technology, such as RAD-seq (Baird *et al.* 2008) and subsequent variants, have also been effective in studying complex traits such as disease resistance in aquaculture species, and testing genomic selection (Vallejo *et al.* 2016; Barría *et al.* 2018; Palaiokostas *et al.* 2018b; Aslam *et al.* 2018). Disease resistance is particularly amenable to genomic selection, because typically it is not possible to record on selection candidates themselves (Yáñez *et al.* 2014), and is typically measured on their close relatives (e.g. full siblings) in aquaculture breeding programmes (Gjedrem and Rye, 2016). While effective, the limitations of current genomic selection methods in aquaculture include (i) that the genotyping is typically expensive, partially due to the high-density marker genotyping, and (ii) the accuracy of prediction drops rapidly when the genetic relationship between the training and validation populations decreases (e.g. Tsai et al., 2016).

Family-based breeding programmes are at a formative stage in common carp, including a programme focused on the Amur mirror carp breed in Europe (Prchal et al., 2018a,b), where improvement of disease resistance is a major breeding goal. The main aim of the current study was to investigate the potential of GS to predict host resistance to KHVD in common carp using genome-wide SNP markers generated by RAD sequencing. An additional aim was to investigate the importance of SNP marker density in genomic prediction accuracy, with a view to future low-density SNP panels for cost-effective genomic selection. Finally, the impact of genetic relationship between the training and validation sets was assessed by comparing prediction accuracy in groups of closely and distantly related fish.

## Materials and Methods

### Population origin and disease challenge

The origin of the samples and the details of the disease challenge experiment have been fully described previously (Palaiokostas *et al* 2018a). In brief, the study was performed on a population of Amur mirror carp that was created at the University of South Bohemia in České Budějovice, Czech Republic in May 2014 using an artificial insemination method (Vandeputte *et al.* 2004). The population was the result of four factorial crosses of five dams x ten sires (20 dams and 40 sires in total). A cohabitation KHV challenge was performed on randomly sampled progeny of these crosses. Mortality of individual fish was recorded for a period of 35 days post infection (dpi), by which stage the mortality level had returned to baseline. In total, phenotypic records regarding survival/mortality were documented for 1,425 animals. Presence of KHV in a sample of dead fish (n = 100) was confirmed by PCR according to guidelines by the Centre for Environment, Fisheries & Aquaculture Science, UK (Cefas) (Pokorova *et al.* 2010). The entire experiment was conducted in accordance with the law on the protection of animals against cruelty (Act no. 246/1992 Coll. of the Czech Republic) upon its approval by Institutional Animal Care and Use Committee (IACUC).

### RAD Sequencing and Parentage assignment

The RAD library preparation protocol followed the methodology originally described in Baird *et al.* (2008), presented in detail in Palaiokostas *et al* (2018a). In brief, RAD libraries were sequenced by BMR Genomics (Padova, Italy) in fourteen lanes of an Illumina NextSeq 500, using 75 base paired-end reads (v2 chemistry). Sequenced reads were aligned to the common carp reference genome assembly version *GCA_000951615.2* (Xu *et al.* 2014) using bowtie2 (Langmead and Salzberg 2012). Uniquely aligned reads were retained for downstream analysis. The aligned reads were sorted into RAD loci and SNPs were identified from both P1 and P2 reads using the Stacks software v2.0 (Catchen *et al.* 2011). Opposed to our previous study (Palaiokostas *et al* 2018a) variant calling in Stacks v2.0 and above utilizes information from both P1 and P2 ends, while prior versions were using only P1 ends. SNPs were detected using gstacks *(--var-alpha 0.001 –gt-alpha 0.001 –min-mapq 40).* Only single SNPs from each individual RAD locus where considered for downstream analysis to minimize the possibility of genotypic errors. SNPs with minor allele frequency (MAF) below 0.05, greater than 25 *%* missing data were discarded. The R package hsphase (Ferdosi *et al.* 2014) was used for parentage assignment allowing for a maximum genotyping error of 2 %. The aligned reads in the format of bam files were deposited in the National Centre for Biotechnology Information (NCBI) repository under project ID PRJNA414021.

### Genomic prediction models

Overall binary survival (0 = dead, 1 = alive) was used as the phenotype to assess the potential of genomic selection for improved resistance to KHVD in common carp. Several commonly used genomic selection models were tested on the data using the R package BGLR for binary traits (Pérez and de los Campos 2014): specifically rrBLUP, BayesA, BayesB (Meuwissen *et al.* 2001) and BayesC (Habier *et al.* 2011). In addition, pedigree-based BLUP (Henderson 1975) was evaluated using the same software. The general form of the fitted models was:

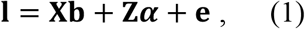

where **1** is the vector of latent variables, **b** is the vector of the fixed effects (intercept, standard length), **X** is the incidence matrix relating phenotypes with the fixed effects, **Z** the incidence matrix relating the underlying liability with the genotypes and ***α*** the vector of SNP effects using the corresponding prior distribution for each of the aforementioned Bayesian models. The parameters of each model were estimated by Markov chain Monte Carlo (MCMC) using Gibbs sampling (110,000 iterations; burn-in: 10,000; thin: 10). Convergence of the resulting posterior distributions was assessed both visually (inspecting the resulting MCMC plots) and analytically using the R package coda v0.19-1 software (Plummer *et al.* 2006).

### Prediction metrics for KHVD resistance

The prediction performance of the utilized models was tested using the following metrics:

- Accuracy
- Receiver operator characteristic (ROC) curves

The prediction accuracy was approximated as:

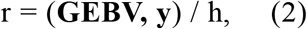

where **y** is the vector of recorded phenotypes, (G)EBV is the (genomic) estimated breeding values and h is the square root of the heritability (*h*^2^ = 0.50 using the genomic relationship matrix as described in Palaiokostas *et al* 2018a).

ROC curves were used to assess the efficacy of classifying the animals as resistant or susceptible, using either the pedigree-or the genomic-based models. The area under the curve (AUC) metric (Hanley and McNeil 1982; Wray *et al.* 2010) was used to interpret the performance of the genomic prediction models, with values of 1 representing the perfect classifier.

### Genomic prediction with varying SNP densities

Genomic prediction models were applied using datasets of varying SNP density <u>using either MAF or linkage disequilibrium (LD) values as thresholds for filtering. In particular</u>, to obtain the reduced density SNP panels for genomic prediction, a strategy of retaining SNPs surpassing a sequentially increased MAF threshold was applied, as described in Robledo *et al.* (2018). These MAF thresholds were 0.1 (3,993 SNPs), 0.25 (1,619 SNPs) and 0.35 (802 SNPs).

In addition, reduced density SNP datasets were obtained by applying filtering based on LD values. LD amongst SNP pairs was calculated using SNPrune (Calus and Vandenplas 2018). Thereafter, only SNP pairs below a sequentially increased LD value were retained. The LD thresholds were 0.15 (1,006 SNPs), 0.25 (2,895 SNPs), 0.35 (5,118 SNPs).

Five-fold cross-validation was performed for all the density varying SNP datasets in order to test the efficiency of correctly classifying animals in the validation set as resistant or susceptible. The dataset was randomly split into sequential training (n = 1008) and validation sets (n = 251). The number of resistant and susceptible animals in each validation set was proportional to the overall survival of the challenged population. In the validation sets, the phenotypes of the animals were masked, and their (genomic) estimated breeding values – (G)EBV – were estimated based on the prediction model derived from the training set. This cross-validation procedure was repeated five times to minimize potential bias.

### Testing the impact of genetic relationship on genomic prediction

Four different scenarios were tested for evaluating the impact of genetic relationships between training and validation sets. In scenario 1 (S1), the formation of training and validation sets required the existence of full-siblings in both sets for each family. For scenario 2 (S2) the formation of validation and training sets allowed the existence of only half siblings between the two sets (and no full siblings). Both in S1 and S2 the cross validation procedure was repeated five times in order to reduce potential bias, while the size of the validation set was 290 animals on each replicate. In scenario (S3) the genomic prediction models were tested by sequentially assigning each of the four factorial crosses (mean = 315 animals; sd = 81 animals) as a validation set, using the remaining three as a training set. This approach resulted in relatively unrelated training and validation sets, since it avoided the inclusion of full / half sibs in both the training and the validation sets. The genomic prediction models were tested on the dataset comprised of the full SNP data. Since pedigree information was not available for prior generations, pBLUP could not be used for obtaining meaningful predictions across the factorial cross groups. Finally, a scenario 4 (S4) was performed as control where no restrictions were applied in the formation of training and validation sets (i.e. they were taken at random). Cross validation in S4 was performed five times with the size of the validation sets being set to 290 animals. The S4 scenario was in fact similar with the approaches tested in the previous section regarding varying SNP densities with the only difference being the size of the validation set. The full SNP dataset was used for all the tested scenarios.

## Results

### Disease challenge

The mean weight of the genotyped carp juveniles was 16.3 g (SD 4.6) and the mean standard length (SL) was 77 mm (SD 7.1). Mortalities began at 12 dpi reaching a maximum daily rate between 21 and 24 dpi (98 – 130 mortalities per day) decreasing thereafter (File S2). Observed mortalities displaying typical KHVD symptoms (weakness, lethargy, loss of equilibrium, erratic swimming, sunken eyes, excessive mucous production, discoloration, and hemorrhagic lesions on the skin and gills).

### RAD Sequencing and Parentage assignment

2.8 billion paired-end reads were uniquely aligned to the common carp genome assembly (GenBank assembly accession *GCA_000951615.2*). Approximately 5 % of those reads had a mapping quality below 40 and were discarded. In total 397,047 putative RAD loci were identified with a mean coverage of 21X (SD = 7.6, min = 1.3X, max = 58.5X). 15,615 SNPs found in more than 75% of the genotyped animals and with a MAF above 0.05 were retained for downstream analysis (File S1).

The carp progeny were assigned to unique parental pairs allowing for a maximum genotypic error rate of 2 %. In total 1,259 offspring were uniquely assigned (File S3), comprising 195 full-sib families (40 sires, 20 dams) ranging from 1 to 21 animals per family with a mean size of 6 (SD 4). The individual dam contribution to the population ranged from 9 to 99 animals with a mean of 61 (SD 23), while the sire contribution ranged from 7 to 53 animals with a mean of 30 (SD 12). In addition, the mean weight and length per full-sib family were approximately 16 (SD 2.8) g and 76 (SD 4.5) mm respectively. Finally, mean survival per full sib family was 34 % (Figure 1).

**Figure 1.**
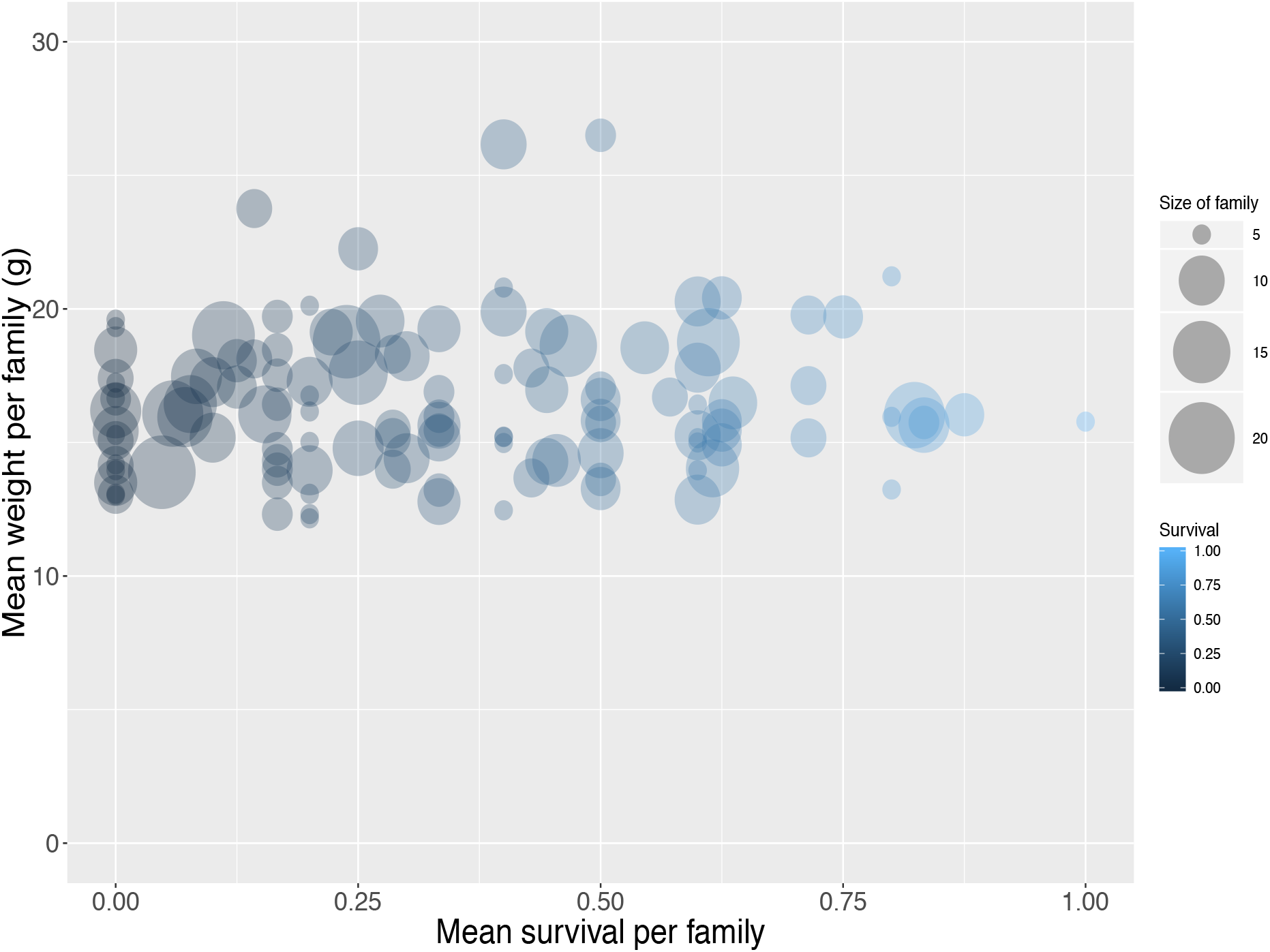
Mean weight and survival for assigned full-sib families.

### Impact of SNP density on genomic prediction

Datasets of varying genotyping density were comprised of 15,615 SNPs (D1; full dataset; File S4) and in the case of MAF as the filtering criterion of 3,993 (D2; MAF 0.1), 1,619 (D3; MAF 0.25) and 802 (D4; MAF 0.35) SNPs. The accuracy of genomic prediction of breeding values was assessed and compared to prediction using a pedigree-based approach. Prediction accuracy with pBLUP was 0.49, compared to 0.53 – 0.54 for the genomic prediction models applied using D1 (Table 1). Prediction accuracies for D2 ranged between 0.52 – 0.53, while in the case of D3 and D4 prediction accuracy for all genomic models was 0.49 and 0.46 respectively (Figure 2a). Following estimation of ROC curves, the genomic models for D1 had a maximum AUC estimate of 0.74 as opposed to 0.71 using pBLUP. AUC for D2 was 0.73 for all genomic models. In the case of D3 and D4 the AUC for all genomic models was 0.71 and 0.70 respectively.

**Figure 2.**
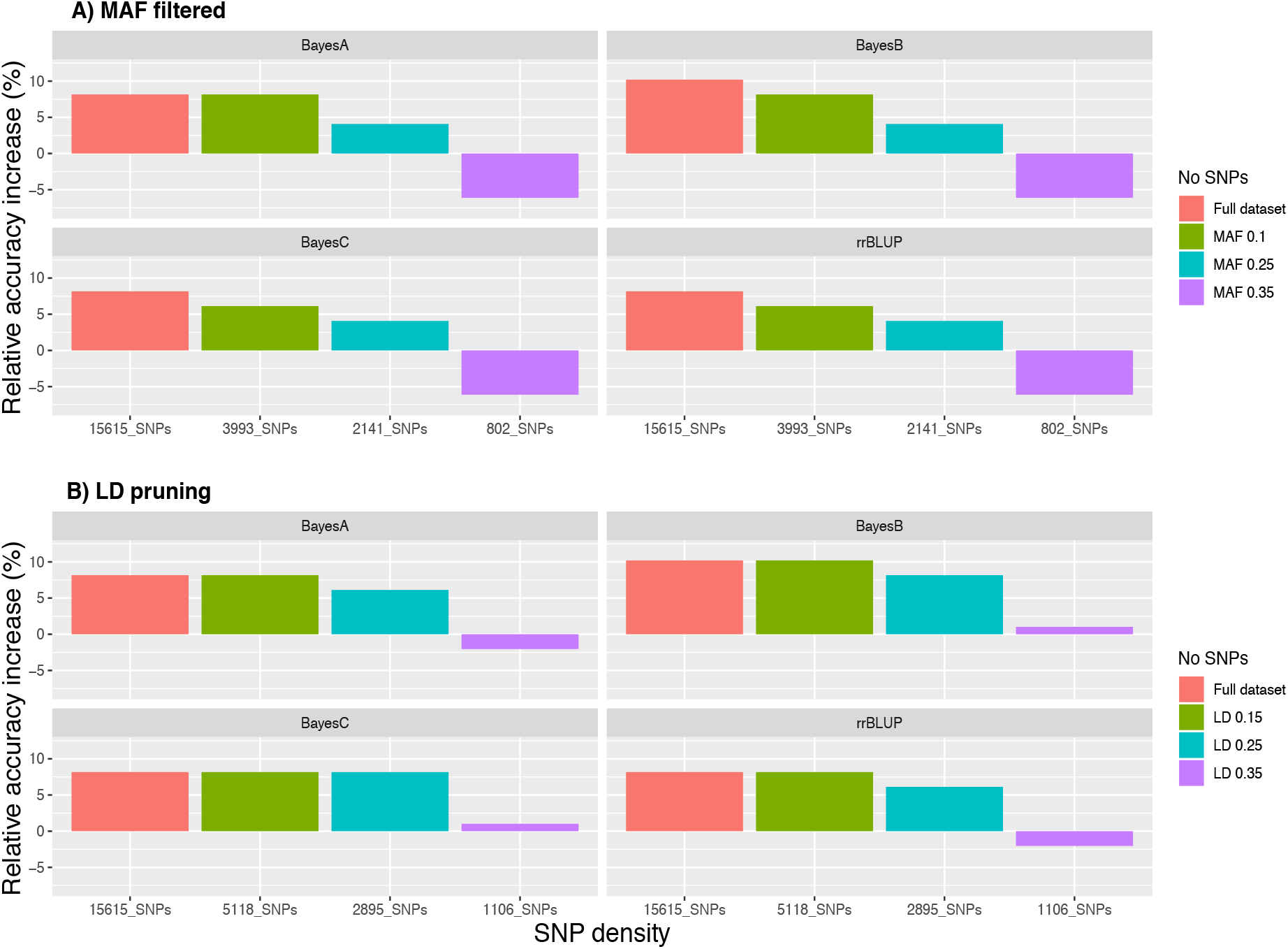
Relative accuracy of genomic prediction models compared to pedigree BLUP for varying SNP densities

**Table 1:** Mean survival accuracy for D1^1^ (5-fold cross validation; 5 replicates)

| Reps | pBLUP | rrBLUP | BayesA | BayesB | BayesC |
| --- | --- | --- | --- | --- | --- |
| 1<sup>st</sup> | 0.47 | 0.51 | 0.52 | 0.52 | 0.52 |
| 2<sup>nd</sup> | 0.49 | 0.54 | 0.55 | 0.55 | 0.54 |
| 3<sup>rd</sup> | 0.48 | 0.52 | 0.53 | 0.54 | 0.53 |
| 4<sup>th</sup> | 0.50 | 0.54 | 0.54 | 0.55 | 0.55 |
| 5<sup>th</sup> | 0.49 | 0.52 | 0.53 | 0.54 | 0.52 |
| **Mean** | **0.49** | **0.53** | **0.53** | **0.54** | **0.53** |

<sup>1</sup>15,615 SNPs

Regarding the reduced density SNP datasets obtained using LD pruning, the number of SNPs in the sets with the LD thresholds of 0.15, 0.25, and 0.35 were 1,006 (LD1), 2,895 (LD2) and 5,118 (LD3) respectively. The genomic prediction accuracy obtained for LD1 was very slightly higher than pBLUP using the BayesB and BayesC models (< 1% increase), while the AUC was the same. In the case of rrBLUP and BayesA for the same SNP dataset the estimates were 2 % and 1 % lower compared to pBLUP for accuracy and AUC respectively. Using datasets of higher SNP density resulted in the increase of both the accuracy and the AUC metrics as observed previously for the reduced density datasets filtered by MAF. In particular, accuracy for LD2 and LD3 ranged between 0.52 – 0.54 and AUC between 0.72 and 0.74 (Figure 2b), which were very similar to the accuracy and AUC values obtained for the full SNP dataset (15,615 SNPs).

### Impact of genetic relationship on genomic prediction

For the scenario S1, where all animals in the validation set had full sibs in the training set the genomic prediction accuracy was approximately 0.56, which was marginally higher (~ 4% increase) than the random allocation of animals into training and validation sets described above. In S2 where the design of the validation set allowed the inclusion of only corresponding half sibs in the training and validation set, the genomic prediction accuracy fell to ~ 0.53. In S3 where the training and validation sets were set up to correspond to separate factorial crosses, the mean accuracy for the genomic models was markedly lower, and ranged between 0.16 – 0.20. Finally, in the scenario where training and validation sets were set up without posing any restrictions estimated, such that close relatives are likely to be included in both sets, accuracy ranged between 0.52 – 0.54 for the genomic prediction models and 0.49 for pBLUP (Table 2).

**Table 2.** Prediction metrics for varying genetic relationships in the validation set.

| Scenario | pBLUP |  | rrBLUP |  | BayesA |  | BayesB |  | BayesC |  |
| --- | --- | --- | --- | --- | --- | --- | --- | --- | --- | --- |
|  | Acc<sup>1</sup> | AUC<sup>2</sup> | Acc<sup>1</sup> | AUC<sup>2</sup> | Acc<sup>1</sup> | AUC<sup>2</sup> | Acc<sup>1</sup> | AUC<sup>2</sup> | Acc<sup>1</sup> | AUC<sup>2</sup> |
| S1 | 0.52 | 0.72 | 0.56 | 0.74 | 0.57 | 0.74 | 0.57 | 0.74 | 0.56 | 0.74 |
| S2 | 0.45 | 0.69 | 0.53 | 0.72 | 0.52 | 0.72 | 0.52 | 0.72 | 0.52 | 0.72 |
| S3 | N/A | N/A | 0.17 | 0.57 | 0.16 | 0.57 | 0.18 | 0.58 | 0.20 | 0.57 |
| S4 | 0.49 | 0.71 | 0.52 | 0.72 | 0.53 | 0.74 | 0.54 | 0.74 | 0.52 | 0.72 |

<sup>1</sup>Accuracy; <sup>2</sup>Area under curve. Genetic relationships for the 4 tested scenarios were for S1: Full sibs on the training set for all animals of the validation set (n=290), for S2: Half sibs on the training set for all animals in the validation set (n=290), for S3: cross validation performed for each of the breeding cross. No full/half sibs on the training set for any of the animals in the validation set (n=315) and for S4: No restrictions applied (n=290) for genetic relationship between training and validation set.

The obtained AUC values from the ROC curves were 0.74 (BayesB; Figure 3) and 0.72 for S1 and S2 for the genomic prediction models, while the corresponding AUC values from pBLUP were 0.72 and 0.69 respectively. For S3 the estimated AUC values for the genomic models were again substantially lower and ranged between 0.57 – 0.58. In S4, where no restrictions were applied regarding the inclusion of full/half sibs on both training and validation sets, the AUC values were between 0.72 – 0.74, comparable to S1 and S2 (Table 2).

**Figure 3.**
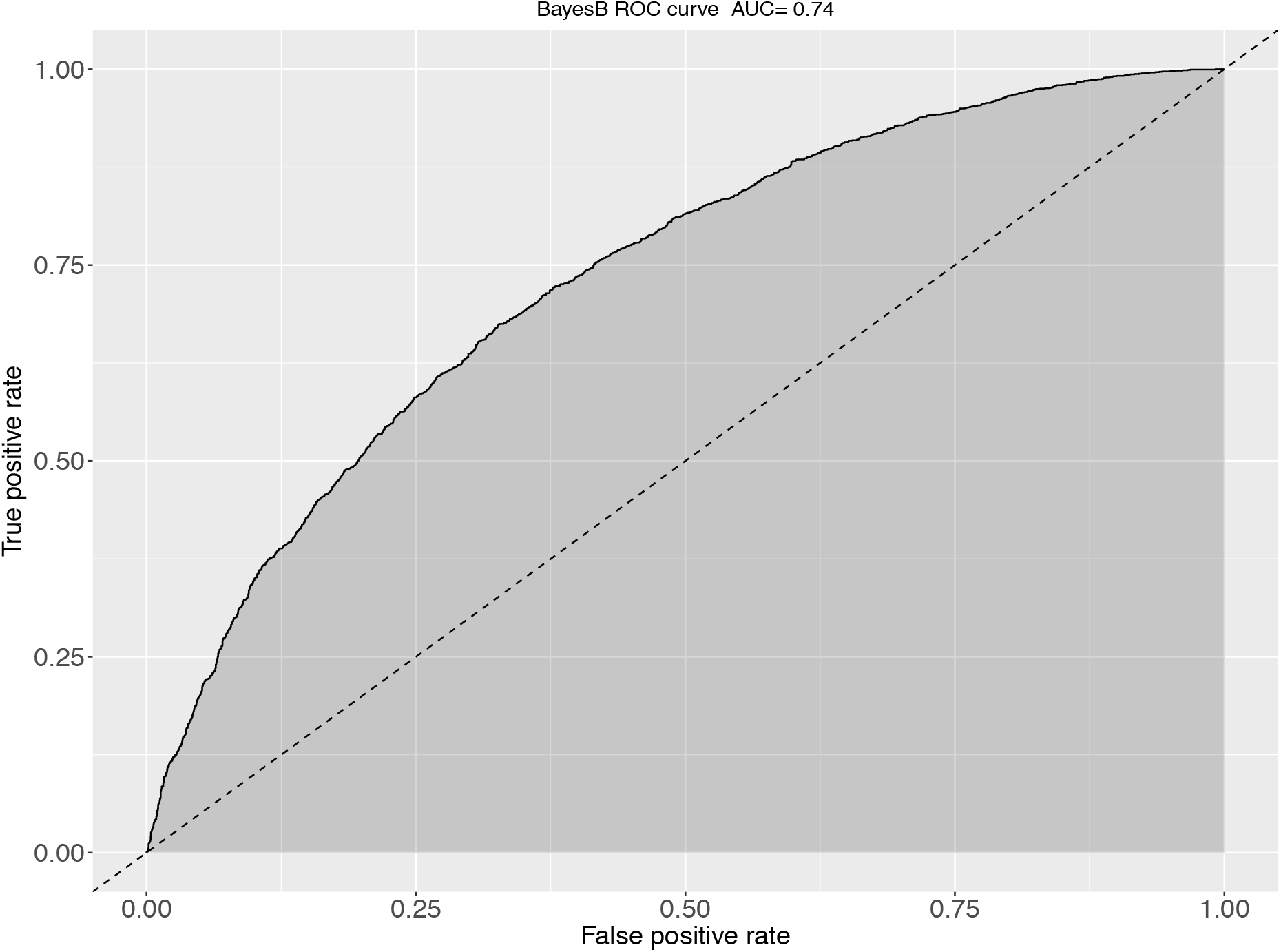
The ROC curve and corresponding AUC metrics for BayesB-based predictions of KHV survival. The plot was obtained from aggregation of a 5-fold cross validation scheme when full sibs existed in the training set for every animal of the validation set.

## Discussion

In the current study, genotyping by sequencing was applied to study genomic prediction of resistance to KHVD in carp, including testing the impact of SNP marker density and genetic relationship between training and validation sets. While genomic data in the form of genetic markers can be a valuable addition to selective breeding for disease resistance, the methods of applying the data depend on the underlying genetic architecture of the trait. In the case of major QTL such as resistance to Infectious Pancreatic Necrosis in salmon (Houston et al. 2008; Moen et al. 2009), it may be most effective to use QTL-targeted marker-assisted selection, and in the case of polygenic traits genomic selection is likely to be preferable. In our previous study we identified a QTL associated with KHVD resistance in common carp located on chromosome 33 (Palaiokostas *et al.* 2018a). However, this QTL accounted for approximately 7 % of the genetic variation in the trait, highlighting that multiple additional loci are involved. Further, using genomic prediction models that incorporate variable selection – i.e. allow for the existence of QTL of large effect – did not result in significant improvement in prediction accuracy compared to ridge regression BLUP, which supports the involvement of many genomic regions in the trait (Meuwissen *et al.* 2001; Kizilkaya *et al.* 2010; Habier *et al.* 2013)

Since genotyping cost is generally related to SNP marker density, determining the lowest SNP density that retains maximum genomic prediction accuracy is a logical goal. In the current study, reducing SNP density from 15,615 to 2,895 resulted in minor decreases in prediction accuracy, with 1,000 – 1,600 SNPs giving approximately the same accuracy as pBLUP. Furthermore, the LD-pruned dataset of approximately 5,000 SNPs resulted in the same prediction accuracy performance as the full dataset (15,615 SNPs). A more drastic impact of genetic relationship between training and validation sets on prediction accuracy was observed. The highest prediction efficiency was observed in scenario S1 where animals in the validation set had full siblings in the training set. Prediction efficiency decreased 6 – 8 % in the scenario allowing for only the inclusion of half-siblings (and no full siblings) in the training and validation sets but was still comparable to the results when the sets were established at random. Interestingly, the impact of the lower genetic relationships on pBLUP accuracy was greater, and it dropped by approximately 16 % between S1 and S2. This may indicate that genomic prediction models have the potential to utilize distant relationships compared to pBLUP, especially in the current set up where there was only a two generation pedigree. Furthermore, when the training set comprised three of the factorial cross groups and the validation set comprised the fourth, thus resulting in no shared full/half sibs between the two sets, the accuracy dropped massively to 0.16 – 0.17 (15,615 SNPs). The decrease in prediction accuracy with more distant relationships is to be expected, thus close relationships between training and validation sets is a necessary prerequisite for successfully implementing GS (Meuwissen *et al.* 2013), and it highlights the importance of obtaining genotype and phenotype records on close relatives of selection candidates in future carp breeding programmes using genomic (and pedigree) selection.

Testing genomic prediction on binary traits such as survival, presents a challenge to define a suitable test metric for selecting the best performing model, especially when survival deviates significantly from 50 %. Solely relying on correlation derived accuracy for model assessment in this case could result in suboptimal selection decisions. Suitable metrics for evaluating prediction efficiency in binary traits and thus selecting the best performing models for estimating breeding values include the AUC from ROC curves.

The AUC values provide a commonly used metric for assessing the prediction efficacy of binary classifiers, taking into consideration both the rate of false positives and false negatives with values of one suggesting 100% successful classification. This approach has been routinely used to test the efficacy of prediction models in disease resistance studies both in humans (Wray *et al.* 2010), livestock (Tsairidou *et al.* 2014) and aquaculture (Palaiokostas *et al.* 2018c) amongst others. In the current study, genomic prediction using the marker density scenarios of ~ 3,000 SNPs and above resulted in a slight improvement (~ 4%) of AUC compared to pBLUP. Performing predictions using approximately 1,000 SNPs resulted in the same AUC value (0.71) as pBLUP, while when using approximately 800 SNP the estimated AUC value was 0.70 which is slightly inferior. A gradual decrease was observed regarding the estimated AUC values for the scenarios of varying genetic relationship as was also the case for the prediction accuracy metric. As expected highest values were obtained in the scenario of highest relationships between training and validation sets (S1). Most striking effect of the impact of genetic relationships between the above sets, however was observed in the scenario where the training and validation sets were set up to be most distantly related, where the estimated AUC values ranged between 0.56 to 0.57, which are substantially lower than all other tested scenarios, but still useful.

In summary the results from the current study demonstrate that GS was more efficient than pBLUP in predicting for KHVD resistant carp. The consistency of improvement in prediction accuracy versus pedigree-based accuracy across multiple scenarios highlights flexibility and robustness to different approaches, and it may allow circumvention of limitations posed by incomplete pedigree records. Of major importance is the fact that relatively low density marker panels could be of value for genomic prediction without loss of accuracy. However, close relationships between training and validation sets are key, with substantial loss of prediction accuracy in the scenario where the sets were relatively unrelated. Pedigree-based prediction was also efficient in scenarios with recorded relationships between training and validation sets, possibly partly because KHVD resistance is a high-heritable trait (h^2^ = 0.5 – 0.79), but genetic markers were required to assign the pedigree in the factorial crosses. Future studies testing the efficiency of single-step BLUP approaches (Aguilar *et al.* 2011; Legarra *et al.* 2014) could potentially prove beneficial by allowing genomic predictions based on larger datasets (only a portion of the dataset would be genotyped, thus reducing costs). Overall our results help inform the use of genetic markers in carp breeding to enable improvement of disease resistance, with downstream benefits of helping prevent KHVD outbreaks in carp aquaculture.

## Author contributions

TV, MK, MP, VP, RH conceived the study, contributed to designing the experimental structure. TV, DP, LP carried out the challenge experiment. CP carried out DNA extractions, RAD library preparation and sequence data processing. CP and RH carried out parentage assignment and the quantitative genetic analyses. All authors contributed to drafting the manuscript.

## ACKNOWLEDGEMENTS

The authors are supported by funding from the European Union’s Seventh Framework Programme (FP7 2007-2013) under grant agreement no. 613611 (FISHBOOST). CP and RH gratefully acknowledge Institute Strategic Funding Grants to The Roslin Institute (Grant Nos. BBS/E/D/20002172, BBS/E/D/30002275, and BBS/E/D/10002070); MK, MP were also supported by project, Biodiverzita (CZ.02.1.01/0.0/0.0/16_025/0007370), VP was also supported by project PROFISH (CZ.02.1.01/0.0/0.0/16_019/0000869) both under the Ministry of Education, Youth and Sports of the Czech Republic; TV, DP and LP were also supported by Ministry of Agriculture of the Czech Republic (project MZE-RO 0518).

